# Unravelling the architecture of Major Histocompatibility Complex class II haplotypes in rhesus macaques

**DOI:** 10.1101/2024.03.26.586730

**Authors:** Nanine de Groot, Marit van der Wiel, Ngoc Giang Le, Natasja G. de Groot, Jesse Bruijnesteijn, Ronald E. Bontrop

## Abstract

The regions in the genome that encode components of the immune system are often featured by polymorphism, copy number variation and segmental duplications. There is a need to thoroughly characterize these complex regions to gain insight into the impact of genomic diversity on health and disease. Here we resolve the organization of complete major histocompatibility complex (MHC) class II regions in rhesus macaques by using a long-read sequencing strategy (Oxford Nanopore Technologies) in concert with adaptive sampling. In particular, the expansion and contraction of the primate *DRB*-region appears to be a dynamic process that involves the rearrangement of different cassettes of paralogous genes. These chromosomal recombination events are propagated by a conserved pseudogene, *DRB6*, which features the integration of two retroviral elements. In contrast, the *DRA* locus appears to be protected from rearrangements, which may be due to the presence of an adjacently located truncated gene segment, *DRB9*. With our sequencing strategy, the annotation, evolutionary conservation, and potential function of pseudogenes can be reassessed, an aspect that was neglected by most genome studies in primates. Furthermore, our approach facilitates the characterization and refinement of an animal model essential to study human biology and disease.

## Introduction

Complex immune regions, such as those of the Major Histocompatibility Complex (MHC) and the Killer Immunoglobulin-like Receptor (KIR) cluster, have generally been rather poorly annotated, as evidenced by gaps and incorrect assemblies in several whole genome sequencing projects covering various species other than humans (He et al. 2019; Warren et al. 2020; Jayakumar et al. 2021). These regions of the genome often display substantial degrees of allelic polymorphism, in concert with significant levels of copy number variation (CNV) and are therefore notoriously difficult to characterize (Martin et al. 2000; Adams and Parham 2001; Vilches and Parham 2002; Trowsdale and Knight 2013).

For this report, we choose to focus on the characterization of the *MHC* class II region of rhesus macaques (*Macaca mulatta, Mamu*) as this species represents an important animal model for studying various aspects of a large array of human diseases (Dijkman et al. 2019; Haque and Levey 2019; Munster et al. 2020; Yu et al. 2020; Böszörményi et al. 2021). Similar as in humans, the *MHC* class II region of the rhesus macaque controls the expression of three distinct allotypes designated *Mamu-DP*, *-DQ* and *-DR* (Daza-Vamenta et al. 2004). These are dimers composed of an alpha and beta chain. A standardized nomenclature system facilitates the discrimination of the different *MHC* genes and alleles (Marsh et al. 2010; de Groot et al. 2012). In brief, duplications within a gene region are sequentially numbered in order of description, and for non-human primates, when possible, the HLA nomenclature was followed (e.g., *DRB1*, *DRB3*). The first two or three digits designate the lineage number, whereas the digits after the first colon label the allelic variation (e.g., *DRB1*01:01*). A third and a fourth set of digits assign synonymous and intronic variations, respectively. Additional suffixes may be used to indicate expression status. For example, alleles that are not expressed, so called null-alleles, are given the suffix “N”. A workshop (“W”) prefix may be used when the gene or lineage status is currently unclear.

Classical MHC class II molecules act as peptide receptors that bind degraded protein segments derived from extracellular pathogens (Roche and Furuta 2015). The dimers are normally expressed on the cell surface of professional antigen presenting cells such as B cells, macrophages, and dendritic cells (Ting and Trowsdale 2002). In the case of an infection, MHC class II molecules loaded with non-self-peptides may activate and amplify the adaptive immune response. More in detail, MHC class II dependent pathways orchestrate and control antibody production or may provide help in eliminating intracellular infections by assisting cytotoxic T lymphocytes. Peptide loading of classical MHC class II molecules is a complex process facilitated by the non-classical HLA-DM dimer (Mosyak et al. 1998). Its chaperone, HLA-DO, also expressed on the membrane of intracellular vesicles, facilitates the loading of immunodominant peptides on HLA-DM. The four genes, encoding these two non-classical class II dimers, map within the *MHC* class II region and display modest levels of polymorphism (Release 3.54 (2023-10)). In contrast, the genes encoding the classical HLA class II molecules display high levels of allelic variability, and susceptibility or resistance to many infectious and chronic diseases is associated with particular alleles or allotypes (Jones et al. 2006; Matzaraki et al. 2017). The central *MHC* class II region also harbors genes that play an important role in transport and loading of peptides on MHC class I molecules (Klein and Sato 2000; Ritz and Seliger 2001). These latter molecules are encoded in another section of the *HLA* region.

The *HLA-DR* region contains a single *DRA* gene coupled to diverse sets of *DRB* genes. The DRA molecule forms a heterodimer with any of the molecules encoded by functional *DRB* genes present in an individual. These DR-dimers are essential for presenting antigens to immune cells. The *DRB* genes display most diversity of the *MHC* class II cluster, reflected by CNV and the presence of pseudogenes, some of which are shared with other primate species (Doxiadis et al. 2012). The physical order of *Mamu-DRB* genes is mapped for two region configurations by sequencing overlapping BAC clones, but these might represent incorrect assemblies due to technical limitations (Daza-Vamenta et al. 2004). On these region configurations, a relatively high number of pseudogenes are defined that lack orthologous equivalents in humans. For other *Mamu-DR* regions, which were previously inferred from amplicon sequencing in combination with segregation studies or from STR-typing strategies, a genomic map is lacking (Doxiadis et al. 2000; Doxiadis et al. 2013; de Groot et al. 2017; de Groot et al. 2022). Hence, the enigma persists for the biological mechanism that makes the *DRB* region prone to expansion and contraction, as well as for the prevalence of conserved and species-specific pseudogenes.

To date, two independent groups succeeded in the assembly of an entire *MHC* haplotype of a cynomolgus macaque (*Macaca fascularis*) by the application of a PacBio long-read sequencing strategy or by a concerted effort utilizing PacBio and Oxford Nanopore Technologies (ONT) (Hu et al. 2022; Karl et al. 2023). Although these *MHC* sequences are of high quality, the comprehensive characterization strategy resolved only the gene content of a single chromosome. The difficulty to correctly assemble the *MHC* region containing segmentally duplicated genes is also shown by numerous gaps and misassembles in the two available rhesus macaque genomes (He et al. 2019; Warren et al. 2020).

Here, we aim to implement a novel sequencing strategy executed on an ONT platform utilizing adaptive sampling to characterize the entire *MHC* class II region in a relatively large panel of rhesus macaques. By resolving this immune region, we intend to enhance the understanding of the mechanisms propelling its diversification, and in particular that of the *DRB* region, in primate species.

## Results

### Definition and assembly of *MHC* class II haplotypes

In this communication *MHC* class II haplotypes are defined as a unique combination of *Mamu-DP*, *-DQ* and *-DR* alleles segregating on a single chromosome. The *Mamu-DR* region displays substantial CNV regarding the beta chain genes. The genetic make-up of a distinctive combination and number of different *DRB* genes is defined as a region configuration (Doxiadis et al. 2000). In the human population only five different *HLA-DRB* region configurations are encountered (Fig. 1), each of them displaying substantial amount of allelic variation (Spies et al. 1985; Gregersen et al. 1986; Andersson et al. 1987; Doxiadis et al. 2008a). In rhesus macaque populations of different geographic origin, the existence of at least 16 *Mamu-DR* region configurations was inferred (Doxiadis et al. 2013). Only a few of them display limited levels of allelic polymorphism. Hence, the strategy to mount a diverse immune response at the population level by the *DR* locus banks in humans mainly on allelic polymorphism, whereas in the rhesus monkey combinational diversity of different *DRB* genes is favoured.

**Figure 1.**
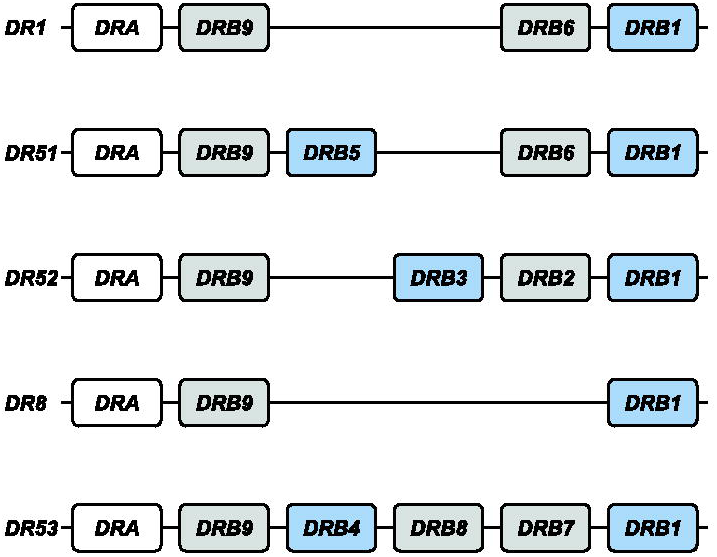
The *HLA-DRB* haplotype groups. In total, 5 major region configurations are defined in humans, containing functional genes (indicated in blue) and pseudogenes (indicated in grey). The *DRA* locus is illustrated as well (white box). All configurations contain a *DRB1* and a *DRB9* entity. The haplotype groups contain 1 or 2 functional genes. The nomenclature of the haplotype groups follows their association with different serotypes. Figure is adapted from (Doxiadis et al. 2008a).

For this communication, we selected 16 heterozygous rhesus macaques, the genomes of which represent 24 unique *Mamu*-class II haplotypes that reflect 17 distinct *DR* region configurations (Suppl. Table S1). Two of these *DR* regions were not documented so far (#17, #18). Our targeted long-read sequencing strategy allowed to assemble and annotate all extended *MHC* class II regions. We choose to report the regions stretching between *Mamu-DRA* and *-DPB2*, as these two genes define fixed locations on the boundaries of the class II region (Fig. 2). As the extended *MHC* class II region is relatively large, the observations made for different subregions are discussed separately in the subsequent paragraphs. The *HLA* region is taken as a reference for comparison.

**Figure 2.**
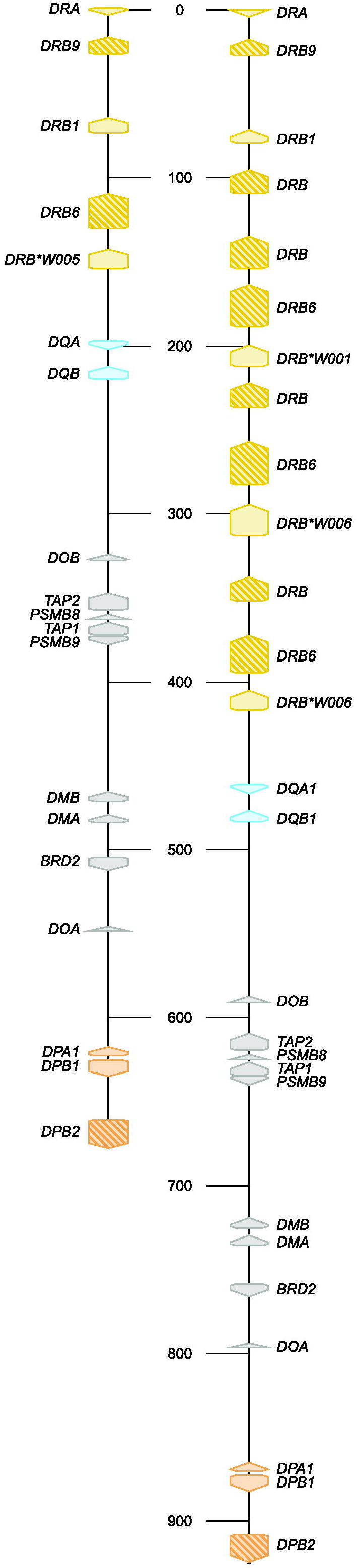
Two illustrative maps of the *MHC* class II region in rhesus macaques. The shortest (A) and longest (B) assembled *MHC* class II haplotypes are depicted at scale and allele level resolution. The genes are indicated by coloured arrows, which indicate the transcription orientation, with the *DR* genes in yellow, the *DQ* genes in blue and the *DP* genes in orange. The other genes encoding products that are involved in peptide processing are depicted with grey arrows. Pseudogenes are indicated with a striped pattern. Allele-level information is provided for all 21 resolved *MHC* class II haplotypes in Supplemental Table S1.

### The *Mamu-DP* region

The human *DP* region encompasses two tandems of genes, namely *HLA-DPA1*, and *-DPB1* and *-DPA2*, and *-DPB2* (Klein and Sato 2000). The latter tandem represents a set of pseudogenes that are characterized by several alterations that render translation into a functional allotype unlikely. The *HLA-DPA1* and *-DPB1* genes are orthologous to *Mamu-DPA1* and *-DPB1*, supported by previous phylogenetic inferences and their fixed location on the chromosome (Fig. 3). In rhesus macaques both genes display polymorphism, which is, however, mainly confined to exon 2 of the *Mamu-DPB1* gene (Slierendregt et al. 1995). The *HLA-DPB1* locus appears to evolve fast due to frequent exchange of polymorphic sequence motifs, a diversifying mechanism that is apparently not active in rhesus macaques (Doxiadis et al. 2001; Otting et al. 2017). The *Mamu-DP* gene content is highly conserved, with only one example of a single configuration that features the genetic relics of a tandem duplication (Suppl. Table S1, region configuration #4). This duplicated set of genes is inactivated, consistent with observed genetic alterations and failure to detect any corresponding transcripts in a previous study (Otting et al. 2017). All 24 haplotypes appear to share the *Mamu-DPB2* pseudogene, which is orthologous to *HLA-DPB2*. The equivalent of *HLA-DPA2* is absent in rhesus macaques and was probably lost during evolution (Fig. 3). The congregate data suggest that the *DPA2-DPB2* tandem arose approximately 30 million years ago in a common ancestor of humans and Old World monkeys and was partly deleted or inactivated before or during evolution of both primate lineages.

**Figure 3.**
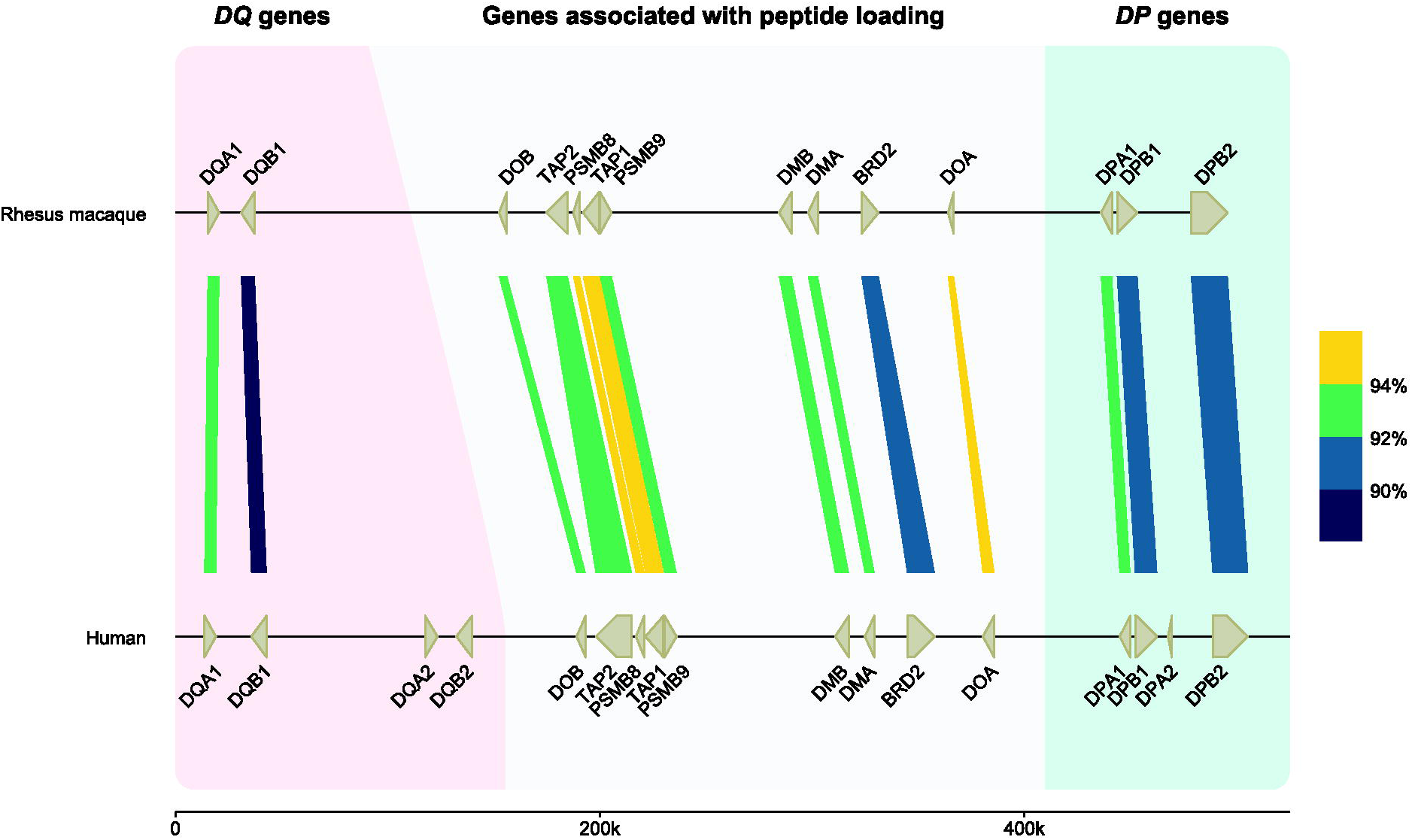
An overview of the similarity between human and rhesus macaques *DQ* and *DP* genes and the members involved in the peptide-loading pathway. On the top lane, a rhesus macaque *MHC* class II region spanning from *DQA* to *DPB2* is illustrated at scale. This high-accuracy region configuration has been resolved using a combination of ONT and PacBio data (total coverage of above 60X) for sequence similarity analysis. On bottom, the human reference genome (GRCh38.p14) is depicted for the equivalent region. The genes and their orientations are indicated by arrows. The colour-coded bars connect homologous human and macaque genes and indicate their sequence similarity percentages that are determined using blastn (see materials and methods). The equivalents of *HLA-DQA2* and - *DQB2* are absent in macaques, as well as a homolog for *HLA-DPA2*. The remaining genes are shared in the two species, some with similarities up to 94%.

### The *Mamu-DQ* genes

The *HLA-DQ* region encodes two sets of genes, namely a *DQA1-DQB1* and *DQA2-DQB2* (Klein and Sato 2000). Our earlier hypothesis about Old World monkeys lacking the equivalents of the *HLA-DQA2* and *-DQB2* genes is substantiated by our latest data, as we found only *Mamu-DQA1* and -*DQB1* genes (Fig. 3) (Bontrop et al. 1999; Doxiadis et al. 2001). In contrast, New World monkeys possess the evolutionary equivalents of both *DQ1* and *DQ2* tandems (Heijmans et al. 2020). In humans, this latter tandem features restricted expression on Langerhans cells and executes an unknown specialized function, whereas in the New World monkey species investigated, it might be expressed on antigen presenting cells and displays limited levels of allelic polymorphism (Bontrop et al. 1999; Lenormand et al. 2012). The duplication of the *DQ* locus probably occurred in the common ancestor of humans, Old World monkeys, and New World monkeys. This was most likely followed by a reversing event that deleted the *DQ2* loci in Old World monkeys. Both genes of the *DQ1* tandem display substantial levels of polymorphism in humans and rhesus monkeys and are organized in a head to tail fashion. Like observed in humans, some *Mamu-DQA1* and *-DQB1* combinations are predominantly linked, as has been reported previously (Otting et al. 2017). Many combinations of alleles are, however, never encountered, which might indicate selection on the potential to form a functional dimer at the cell surface. A similar form of selection has been proposed for *HLA-DP* alleles (Hollenbach et al. 2012).

#### The MHC class I and II peptide loading pathway genes

Positioned between the *Mamu-DQ* and *-DP* regions lies a cluster of genes whose products are involved in loading MHC class I and II molecules with allotype-specific peptides (Fig. 3) (Klein and Sato 2000). The *DOB* and *DOA* genes are separated by *TAP2*, *PSMB8*, *TAP1, PSMB9*, *DMB* and *DMA,* and *BRD2*, respectively. This part of the *HLA* region and its rhesus macaque equivalent appears to be highly conserved, including the transcriptional orientation of the genes.

The *Mamu*-*DMA* and *-DM B* genes display polymorphism, with 9 and 7 alleles documented for these genes, respectively, but almost half of them involve only synonymous mutations (Suppl. Fig. S1). More variable sequences are observed for *Mamu*-*DOA* and -*DOB*, with 13 and 11 different alleles, respectively (Suppl. Fig. S2). The DM- and DO-dimers are essential elements of the MHC class II peptide loading pathway. At this stage it is not yet understood whether allelic polymorphism in these genes has any functional relevance, but it may involve preferential loading of immunodominant peptides in the context of particular MHC class II allotypes that are encoded on the same haplotype. Also, the *TAP1* and *TAP2* genes in rhesus macaques appear to be polymorphic and within the panel at least 12 and 15 different alleles were encountered, respectively (Suppl. Fig. S3). Population studies are needed to sort out whether polymorphic *TAP* genes segregate with certain *MHC* class I region configurations/alleles. It is tempting to speculate that this phenomenon may explain long range linkage disequilibria as has been recorded for other species (Powis et al. 1996; Walker et al. 2011).

### Introducing the *DR* region

The *DR* region comprises a single *DRA* gene in combination with different sets of *DRB* genes. The *Mamu-DRA* locus is highly conserved, similar to its human equivalent, and the different alleles appear to be indicative for particular *DR* region configurations suggesting fixation. The *HLA-DRB9* pseudogene maps directly adjacent to the invariant *DRA* gene (Klein and Sato 2000). In rhesus macaques we recovered *Mamu-DRB9* at the same location and it is dysfunctional as well, due to many sequence inconsistencies. Phylogenetic analysis of the exon 2 data illustrated that *HLA-* and *Mamu-DRB9* are orthologous to each other (Suppl. Fig. S4). Like found in humans, this gene is present on all rhesus macaque region configurations analysed thus far (Suppl. Table S1) and represents an old entity that was inactivated long ago.

Eight other *DRB* genes have been identified in humans, named *HLA-DRB1* to *-DRB8,* which are distributed among 5 distinct region configurations (Fig. 1). In addition to *DRB9*, we encountered orthologous sequences for four of these genes in rhesus macaques, while no matching sequences were identified for the *HLA-DRB7* and *-DRB8* pseudogenes (Fig. 4). The *HLA-DRB2* pseudogene is paralogous to *HLA-DRB6*, originating from a duplication that occurred after speciation of humans and macaques (Doxiadis et al. 2008c). In contrast to *DRB9*, the orthologues *DRB1* (as well as some of its lineages), *DRB5*, and *DRB6* genes do not necessarily share their genomic locations (Fig. 4).

**Figure 4.**
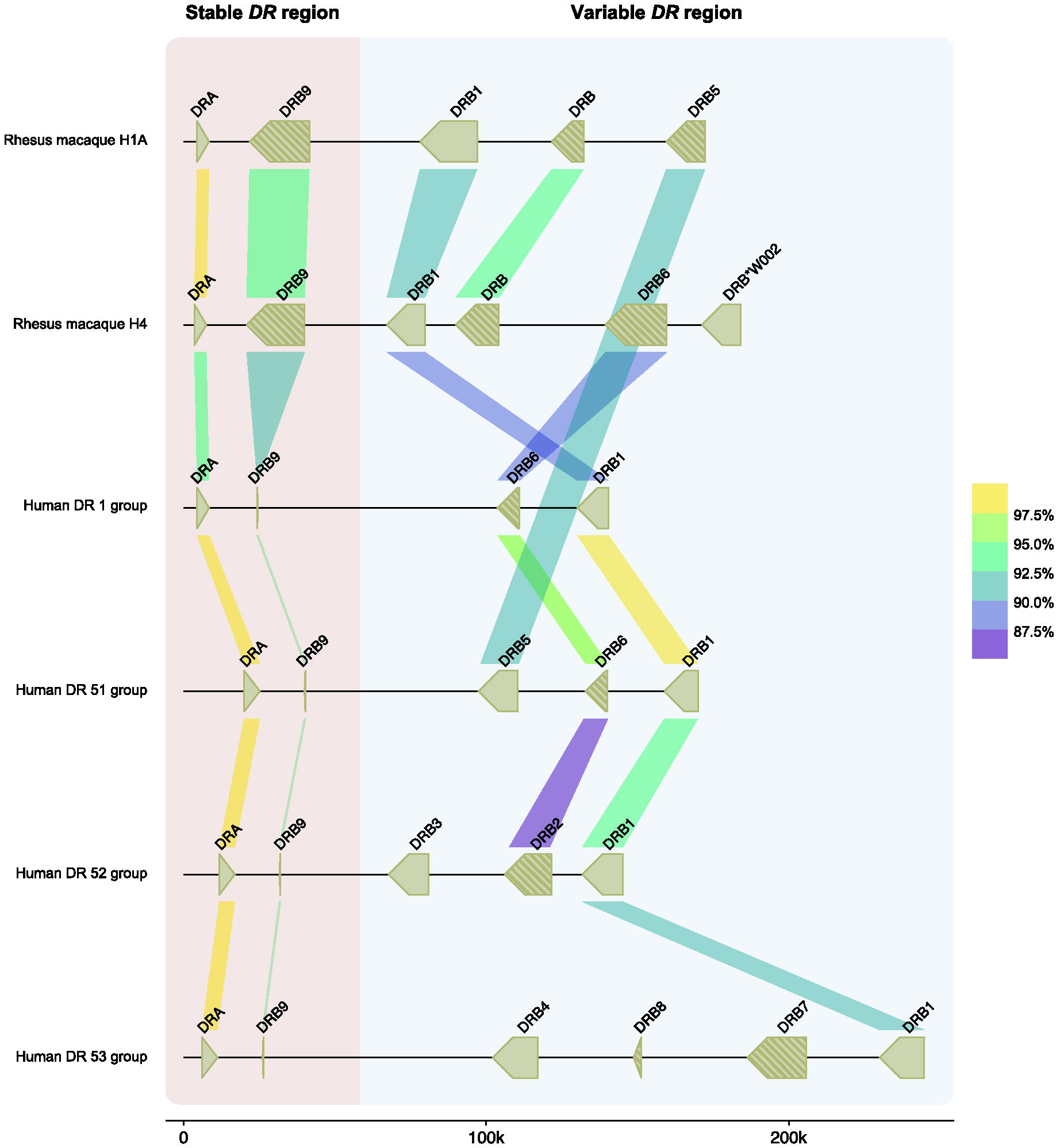
A schematic overview of the similarity between human and rhesus macaque *DR* genes. Two rhesus macaque *DR* region configurations (#1A and 4) are depicted at scale, which are assembled using a combination of ONT and PacBio data (> 60X coverage). For comparison, four human *DR* region configurations, representing the DR1, DR51, DR52, and DR53 group haplotypes, were extracted from the NCBI database (GRCh38.p14: NC_00006, NT_167249.2; OK649233; NT_113891) (Houwaart et al. 2023). The *DR* genes and their orientations are represented by arrows. A striped pattern indicates pseudogenes. The colour-coded bars connect homologous human and macaque genes and indicate their sequence similarity percentages that are determined using blastn (see materials and methods). The *DRA* and *DRB9* loci represent a stable stretch in the *DR* region, whereas a more diverse gene content is encountered for the remaining haplotype. In the selected macaque and human region configurations, four *DRB* genes were determined to be orthologs, with similarities up to 90-93%, whereas the *DRA* gene is even more conserved. The rhesus macaque *MHC* class II genes were originally named based on resemblance of their exon 2 sequences to *HLA* equivalents (de Groot et al. 2012). In the case that such an equivalent was absent in humans a W (workshop) assignment was introduced, for instance for *Mamu-DRB*W002*.

Although some *DRB* genes and lineages are shared, none of the *HLA-* and *Mamu-DR* region configurations are identical in composition (Fig. 1 and Fig. 5), alluding to a far more dynamic type of evolution as compared to *DQ* and *DP* gene regions. The rhesus macaque *MHC* class II haplotypes differ substantially in physical length, which is mainly due to CNV encountered on the different *Mamu-DRB* region configurations (Fig. 5). Up to 11 different *DRB* entities might be present per region configuration, of which 2 or 3 may be functional (Fig. 5; indicated by blue boxes). The smallest *Mamu-DRB* region configuration, with two functional genes (#2), measures 110 kb, whereas the longest with three intact genes (#16) ranges up to 370 kb. Through the examination of our genomic data, we revealed that most haplotypes carry a substantial number of unannotated pseudogenes (Fig. 5; marked in grey boxes). Most of these inactive genes display variation, reflected by a wide array of mutations in addition to truncations. These pseudogenes often consist of segments from functional *DRB* genes that are present in the contemporary population (Suppl. Fig. S5). This suggests that these genes were relatively recently inactivated, probably during a recombination process or afterwards.

**Figure 5.**
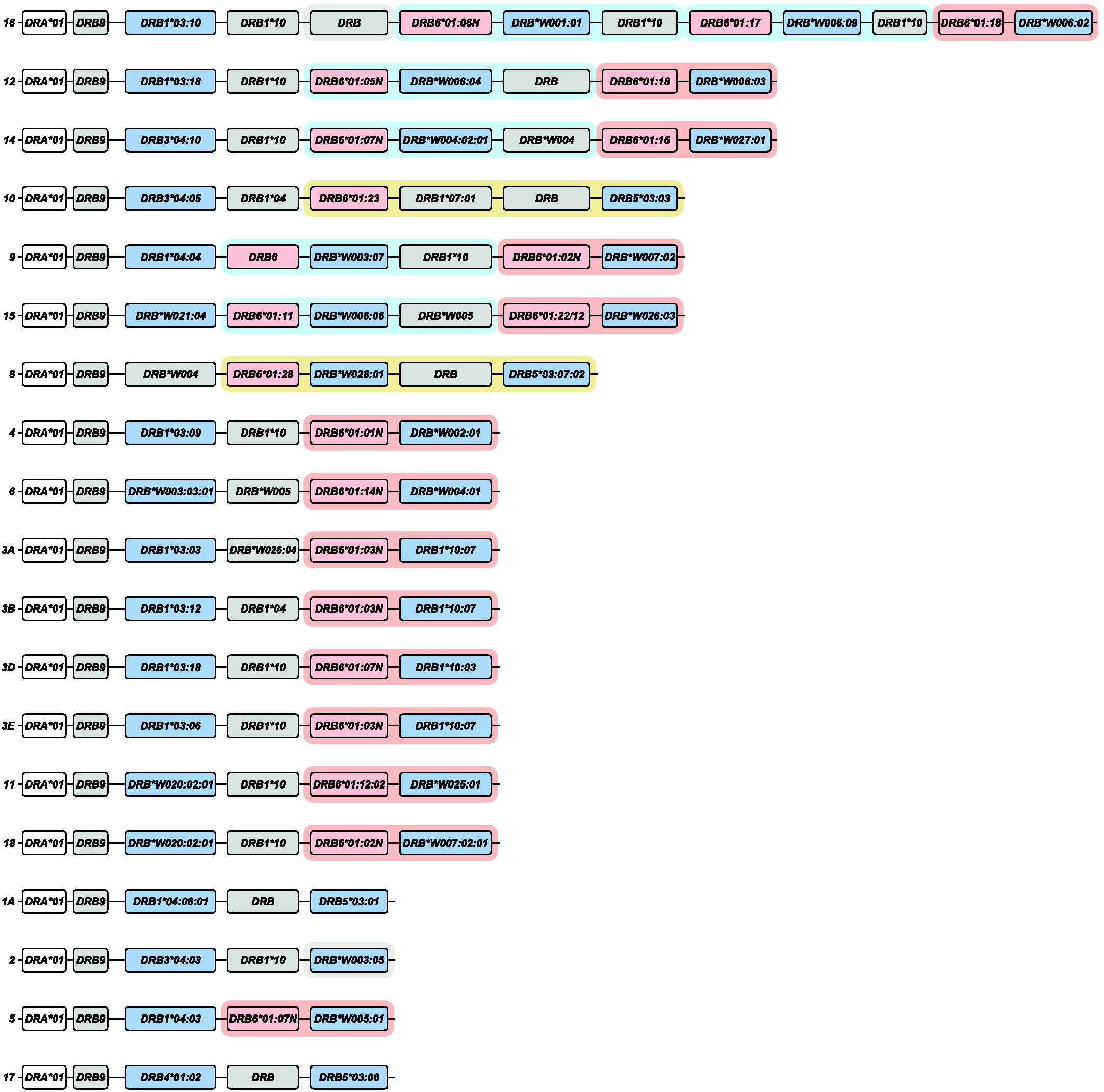
A schematic representation of the composition of distinct rhesus macaque DR region configurations. The different region configurations analysed in this cohort are numbered according to a previous report (Doxiadis et al. 2013). Functional genes are indicated in blue boxes, whereas pseudogenes are in grey. These inactive genes are designated based on the functional allele with which they exhibit at least 95% sequence similarity, whereas inactive entities lacking similarity are denoted as “DRB”. Pseudogene *DRB6* is an exception and is depicted with red boxes. The *DRB9* and *DRA* loci (white boxes) represent a framework that is structurally consistent on every region configuration. The locus equivalent to *HLA-DRB1* is often occupied by a homolog, but other genes are encountered as well. The shadings in four different colours, connecting multiple gene boxes, indicate the different cassettes of paralogous genes. The cassettes are distinguished by their gene content, which is indicated in Figure 6. These cassettes always start with a *DRB6* copy, which propels the region diversification via two instable retroviral elements. Two other genes, a truncated one on region configurations #16 (DRB) and a functional gene on configuration #2 (*DRB*W003:05*), also comprise one of these LTRs and might form a rearranging cassette on their own (depicted with a grey shading).

### *DRB* Region configuration diversity is driven by cassette shuffling

A non-coding stretch that contains repetitive elements separates the genetically stable *DRA-DRB9* framework from a first functional *DRB* gene. Different genes are documented at this locus, including the *Mamu-DRB1*03/04*, -*DRB3*04*, -*\*W003* and *-*W020* lineages, which are neighboured by a truncated pseudogene (Fig. 5). The origin of these truncated genes is complicated, but we detected remnants of *Mamu-DRB1*04*, *-DRB1*10* and *-DRB*W005*. One configuration (#16) contains a second truncated *DRB* remnant, which is not present on any other configuration analysed (Fig. 5). Two functional *DRB* genes located at the position of the *DRB1* locus, *Mamu-DRB1*04:04* and *-DRB*W021*, were not accompanied by a truncated segment, whereas region configuration #8 is an outlier as reflected by the presence of only a pseudogene, *DRB*W004:01*, at this position (Fig. 5).

The next section of the *DR* region is subject to a dynamic expansion and contraction process, evidenced by the shuffling of cassettes containing paralogous genes and gene remnants by recombination events (Figs. 5 and 6). We distinguish three main cassettes based on their gene content and each cassette starts with a *DRB6* gene. This pseudogene is present on approximately 80% of the *Mamu-DR* region configurations analysed (Fig. 5). In addition to this *DRB6* pseudogene, the organization of cassettes I to III comprise a tandem of a functional and a truncated gene, a single functional gene, or a combination of three genes, respectively. Most *DRB* region configurations have one of these cassettes present, whereas five haplotypes have up to three cassettes, which indicates multiple consecutive rearrangement events. Three configurations (#1A, #2 and #17) lack a cassette and might represent stable organizations, which is supported by their relatively high frequencies reported previously (Doxiadis et al. 2013).

**Figure 6.**
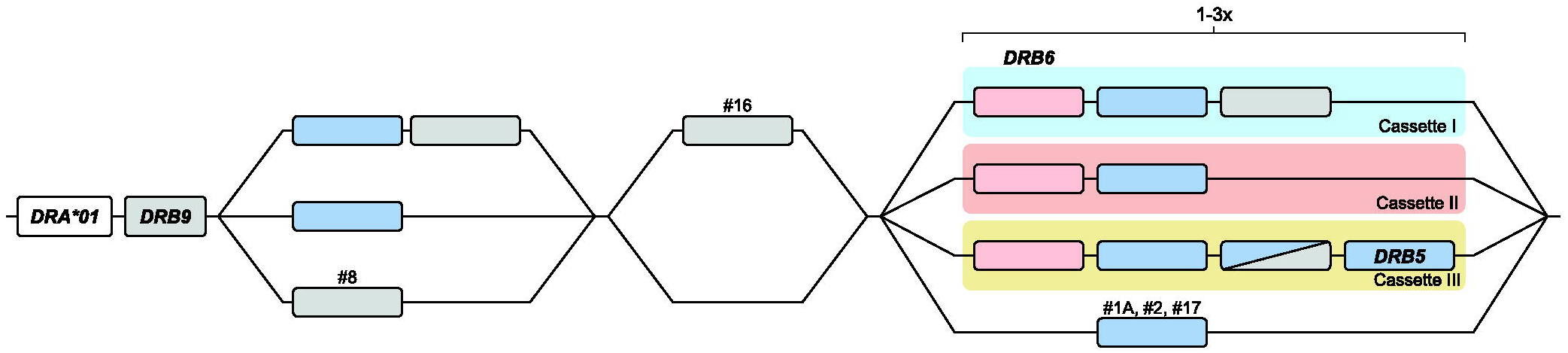
A schematic representation of the building blocks that form the rhesus macaque *DRB* region configurations. All rhesus macaque *DR* haplotypes contain a *DRA* gene, followed by the presence of a *DRB9* segment in close proximity, which consists of exon 2 and its flanking introns. This *DR* framework is separated from the remaining DRB haplotype by a non-coding stretch that varies in length due to the presence of different repetitive elements. The *DRB* region starts with either a functional gene in tandem with a truncated pseudogene or with a single functional gene. Region configuration #8 deviates from this organization, as the first functional gene is absent and instead only a defective gene is identified at this locus. A second exception to the most common organizations is represented by an additional isolated pseudogene at region configuration #16. The remaining part of the *DRB* haplotypes display more variation, with the presence of 1 to 3 cassettes that can contain different paralogous genes (indicated in with coloured shadings). Three region configurations lack a cassette and only have a single functional gene present at this position (#1A, #2, #17).

Region configuration #16 features two copies of cassette I in conjunction with cassette II, representing the longest haplotype. An identical copy of this cassette II is also identified on region configuration #12. Hypothetically, configuration #16 could have originated from configuration #12 by an introduction of cassette I with the functional *DRB*W006:09* gene, substantiating the shuffling of defined cassettes. On region configuration #14 the functional copy of the *DRB*W004* gene in cassette I is phylogenetically highly related to the truncated copy it is segregating with, suggesting a rather recent inactivation during or after its duplication. On other configurations containing cassette I (#9, #12, #15 and #16) any apparent phylogenetic relationship of genes is absent. The functional gene present in cassette I comprise the *DRB*W001*, *-W003*, *-W004* and *DRB*W006* lineage members. These genes are, however, not restricted to these cassettes, as an “isolated” *DRB*W003* lineage member is also found on region configuration #2. This observation may highlight that over long time spans these cassettes are not stable and that uncoupling of particular genes may occur through recombination events within cassettes.

Cassette II comprises a *DRB6* gene family member with a functional gene. The lineages encountered for the functional gene are *Mamu-DRB1*10*, *-DRB*W002*, -*W004, -W005, - W006, -W007*, -*W025*, -*W026* and -*W027*. This tandem cassette might be present alone, characterizing region configurations with two functional genes, or in combination with a cassette I. The longest cassette III is the least frequent one and is only present on region configurations #8 and #10. In both these cassettes III a functional *DRB5* gene is present, which is not associated with any of the other cassettes. Instead, “isolated” copies of a *DRB5* are identified on region configurations that lack the presence of a cassette (#1A and #17). Most likely cassette III originates from a rearrangement of cassette I with a region configuration that lacks a cassette. However, so far, the other expected end-product of this rearrangement, a configuration with only one functional gene and one remnant, has not been identified.

The shuffling of cassettes generates a plastic system and makes it difficult to comprehend the locus or lineage status of the different *DRB* genes. An apparent side effect is the generation of a myriad of pseudogenes and segments thereof.

### Towards understanding the role of the conserved *DRB6* pseudogene

All three cassettes start with the *DRB6* pseudogene, which has an integrated Mouse Mammary Tumour Virus (MMTV) with strong long terminal repeats (LTR) (Mayer et al. 1993). This integration probably inactivated the *DRB6* gene long ago, but the LTR took over the promotor function and drives, if present, transcription of the first exons in humans, chimpanzees, and macaques (Paz-Artal et al. 1996; Fernandez-Soria et al. 1998; Moreno-Pelayo et al. 1999). Characterization of complete genomic regions allowed us to identify a second LTR profile that is associated with *Mamu-DRB6* (Fig. 7). This LTR measures approximately 6 kb and is situated in the 3’ flanking region. A highly similar LTR maps to the 3’ flanking region of *HLA-DRB5*, indicating that a rearrangement has introduced this second repetitive element to *Mamu*-*DRB6* after speciation of humans and macaques. We uncovered one allele, *Mamu-DRB6*01:11* on region configuration #15, that does not contain the second LTR. The combination of two LTRs that in general associate with the macaque *DRB6* gene is likely to facilitate the expansion and contraction of the *DR* region by homologous recombination events. However, the LTR located in the 3’ flanking region might be predominantly involved, as copies of this repetitive element are also encountered in context of two other macaque *DRB* genes. One of these genes is truncated and located in front of the *DRB6*-associated cassettes on region configuration #16. The other is the functional *DRB*W003:05* on region configuration #2. On a similar configuration that was assembled for a rhesus macaque of mixed origin, another *DRB*W003:05* copy was identified featuring the same LTR, which indicates a relatively old integration (Suppl. Table S1). The presence of this element adjacent to the 3’ end of the two *DRB* entities make them potentially mobile cassettes comprising a single gene. This would explain the unregular position of the truncated *DRB* gene on region configuration #16 (Fig. 5). The *DRB*W003:05* gene on configuration #2 shares its putative locus with *DRB5* on two largely similar configurations (#1A and #17), which indicates that the LTR-associated gene might have replaced a *DRB5* gene by chromosomal rearrangement.

**Figure 7.**
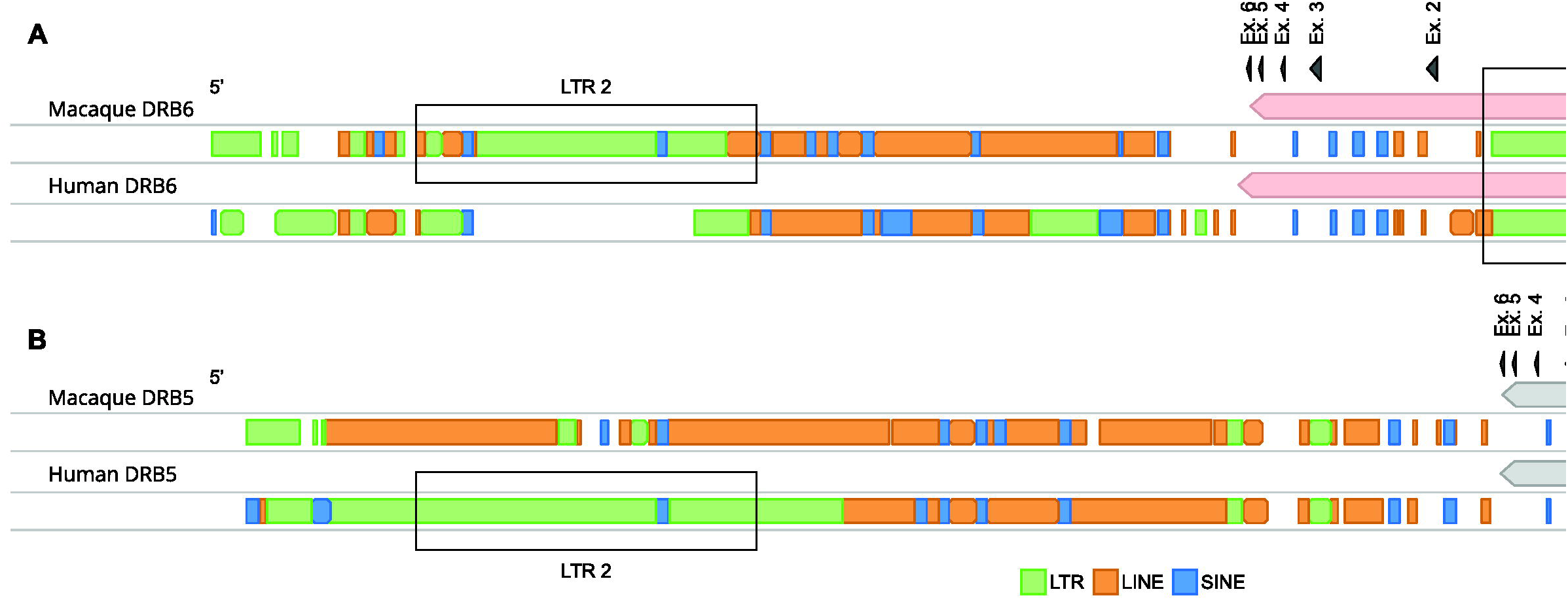
The different repetitive elements associated with *DRB6* and *DRB5* genes. The macaque (top) and human (bottom) *DRB6* (A) and *DRB5* (B) genes and their 3’ flanking regions are illustrated. The exons are depicted as arrows. The repetitive elements, which are determined using RepeatMasker, are given and distinguishes LTRs (green), LINEs (orange) and SINEs (blue). Macaque *DRB6* contains two strong LTR elements, indicated in the boxes; one located in intron 1, whereas the other one is integrated into its 3’ flanking region. This latter LTR originates from a *DRB5* sequence that was present in a shared progenitor.

## Discussion

We demonstrated that the human and rhesus macaque *MHC* class II regions share a similar organization, featured by the presence of the genes encoding the *DR*, *DQ*, and *DP* allotypes. In humans, the *DQ* and *DP* loci are occupied by a tandem of alpha (*A*) and beta (*B*) genes (Fig. 3). Old World monkeys lost the genes equivalent to *HLA-DQA2*, *-B2* and *HLA-DPA2*, whereas relics of *Mamu-DPB2* persist. The *DR* region displays most variation, which is reflected by substantial copy number variation. In humans five *DRB* region configuration have been identified and each of them display high levels of allelic variation (Fig. 1). In contrast, rhesus monkeys feature diverse *DRB* region configurations that exhibit limited allelic variation (Fig. 5). Humans and rhesus macaques do not share a single region configuration, although some of the *DRB* genes and lineages predate their speciation.

Of the nine different human *DRB* genes, only three or four are functional, whereas the remaining genes represent pseudogenes (Doxiadis et al. 2012). The exception is *HLA-DRB4*, which might be active or inactive depending on the allele content of the region configuration. Orthologs of *HLA-DRB1* and -*DRB5* are present in Old World monkeys, suggesting a strong selective pressure to conserve these old functional genes. The DR dimers composed of different functional DRB subunits (*DRB1*, *DRB5*) present largely complementary peptide repertoires to T cells, which might be the reason for the maintenance of both functional genes in several primate species (Scholz et al. 2017). The orthologous *DRB6* and *DRB9* pseudogenes in humans and macaques do not encode functional subunits that are involved in peptide presentation, and their conservation might implicate other important functions.

Pseudogenes are broadly defined as non-functional sequences in the genome that are derived from one or more functional paralogous genes (Balakirev and Ayala 2003). These pseudogenes generally lose their protein-coding potential due to the disruption of promotor or enhancer sequences, or by the accumulation of mutations that cause frameshifts or introduce early stop codons. Also, the integration of retroviral elements may inactivate an existing gene. With the emergence of novel sequencing technologies, more pseudogenes are annotated, and their function have been reassessed. Although the majority indeed represent defective gene copies, a relatively large fraction seems to play an important role in biological processes (Cheetham et al. 2020). For example, pseudogenes have been associated with the production of truncated proteins that have intact functional domains, with the regulation of transcription and translation through the generation of small interfering RNAs (siRNA) and long non-coding RNAs (lncRNAs), with the 3D remodelling of chromatin structures, and with recombination processes, such as gene conversion (Cheetham et al. 2020). These protein-independent functions might explain the conservation of “pseudogenes” that still have biological implications, whereas non-functional defective genes might be released from selective pressure and accumulate variation.

An example of an old pseudogene is *DRB9*. The *HLA-DRB9* remnants consist of a single exon 2, which is flanked by its intron sequences, whereas the remaining gene segments are absent (Gongora et al. 1997). A similar non-coding remnant is present in rhesus macaques, but the original intron-exon organization of the *Mamu*-*DRB9* genes has been remained more intact (Fig. 4). The conservation of exon 2 in *DRB9* genes might associate with its fixed position in close proximity to the shared *DRA* locus. The DRA molecule is essential to generate a functional DR dimer at the cell surface that can present peptides to other immune cells. Local recombination events might be suppressed around this locus to avoid deletion or truncation of an essential element of the dimer. Recombination coldspots are, however, poorly characterized and may involve different factors, such as epigenetic modifications, repetitive elements and chromatin structures (Arnheim et al. 2003). Therefore, it is unclear whether the conserved *DRB9* remnants contribute to this protection by stabilizing the region or that its maintenance is mediated via preserving surrounding sequences.

On the contrary, we suggest that the pseudogene *DRB6* appears to be a hotspot for recombination events in macaques. All three defined cassettes that rearrange in the *DRB* region start with the presence of this truncated pseudogene. The chromosomal instability that is associated with *Mamu-DRB6* is most likely caused by the presence of two LTR elements in and adjacent to the pseudogene (Fig. 7). The first LTR represents an endogenous retroviral element that relates to the MMTV and has been integrated into intron 1 of *DRB6* before the speciation of humans and Old World monkeys (Mayer et al. 1993). In this study, we identified a second LTR in the 3’ flanking region of all *Mamu-DRB6* alleles, except for one copy on region configuration #15. In addition, this LTR element was also identified downstream of a truncated gene (region configuration #16) and of a functional *DRB*W003:05* gene (region configuration #2). These instable retroviral sequences might align during meiosis and initiate homologous chromosomal recombination, explaining the observed reshuffling of cassettes containing a *DRB6* gene. Although details on the precise mechanism are lacking, these kind of LTR-LTR homologous recombination events have been documented before (Lovett 2004; Belshaw et al. 2007; Gu et al. 2008; Thomas et al. 2018). Our hypothesis is substantiated by the peculiar position of the two other gene entities that contain the same LTR on region configurations #2 and 16. These genes may represent a rearranging cassette on its own (Fig. 5). In humans, a highly similar LTR is identified adjacent to the 3’ end of *HLA-DRB5*, which is present on the DR51 group of haplotypes (Fig. 1 and Fig. 7). The absence of a similar retroviral element on any of the other haplotype groups may hamper the initiation of homologous recombination events during meiosis, which might explain the rather fixed set of five *DRB* region configurations in humans. In addition, the association of two LTRs with macaque *DRB6* expands the overall length of a homologous and molecularly instable stretch (Fig. 7), which might enhance the local chromosomal recombination rate. Like in humans, the *DRB5* equivalent in chimpanzees displays a similar LTR in its 3’ flanking region, indicating that after speciation a rearrangement has introduced this retroviral element in the macaque *DRB6* gene. This defective gene is present on approximately 80% of the documented region configurations. New World monkeys, like common marmosets, lack the equivalent of *DRB6* and its associated LTR elements, and have rather simple *DR* regions that do not display CNV (Antunes et al. 1998). These observations shed light on the phenomenon that a pseudogene like *DRB6* has been maintained over long periods of evolutionary time, probably driven by its newly acquired function to promote diversity in the *DR* region of primates.

It has been a puzzle why the *HLA-DRB1*03* (DR52 family) and *\*04* (DR53 family) lineage members positioned at the same locus are differentiated by such large genetic distances. Members of these two lineages are present in macaques as well, indicating that the orthologous *DRB1* lineages predate speciation. Sequence diversity between lineages might be caused by exon 2 shuffling, an ancient exchange of polymorphic gene segments between closely related *DRB* genes. This is evident for the the *HLA-* and *Mamu-DRB3* genes, and most likely also apply to the *DRB4* genes (Doxiadis et al. 2008b). A second possibility is that these gene lineages arose through duplication, initially had paralogous relationships, and the accumulation of point mutations and gene conversion subsequently resulted in their genetic divergence (Gorski et al. 1984). It is likely that the two identified LTRs, associated with *HLA-DRB5* and *-DRB6* and with *Mamu-DRB6*, facilitated recombination events that placed different paralogs in orthologous positions during evolution. Apparently, this diversifying process is still active in macaques, which display a high level of *DR* region diversity. This is exemplified by the large number of genetically different lineages that occupy the putative *DRB1* locus. Its fluidity is illustrated by *Mamu-DRB*W003*, which is present at the supposed *DRB1* locus on region configuration #6, whereas cassette I on configuration #9 features a gene of the same lineage, indicating a paralogous relationship.

Expansion and contraction of the *DRB* region may also have fundamental implications, which could be considered at two different levels. The first one concerns the *MHC* class II repertoire at the individual level. The number of functional genes that encode for polymorphic DRB molecules per region configuration ranges typically from 2 to 3. A single region configuration (#16) forms an exception, encoding four functional DRB molecules. If the number of functional *DRB* genes would increase too much, this might impact the number of T cells that are deleted during thymic selection, and thereby the ability to mount broad levels of CD4-mediated T cell responses (Huseby et al. 2005). We identified inactivated *DRB* entities in the rhesus macaque *DR* region, which do not relate to the *DRB6* pseudogene. Most of these defective genes appear to be paralogous to functional *DRB* genes. In comparison, the *KIR* region is known to evolve relatively fast by recombination events, but inactivated genes are not encountered (Bruijnesteijn et al. 2020; de Groot et al. 2023). This suggest that the diversification of the primate *DRB* and *KIR* immune regions is propelled by different mechanisms. It is noted that especially the region configurations that encode one or two functional DRB molecules are characterized by high frequencies in the rhesus macaque population studied. This might indicate a negative selection on haplotypes that display more functional genes, as an extended DRB repertoire might compromise an efficient T cell population. The second level relates to the population dynamics of *MHC* class II evolution in primates. Diversity of the *DR* region in the human population is largely banking on allelic polymorphism whereas in rhesus macaques natural selection has favoured the presence of unique combinations of *DRB* genes (Doxiadis et al. 2001). Both strategies warrant that individuals within a given human or rhesus macaque population are diverse either by a combinatorial repertoire of *DR* alleles or genes, respectively, thereby minimizing the chance that a single pathogen may exterminate an entire population.

Many of the *MHC* class II genes and their lineages in humans and rhesus macaques are old, predate their speciation and may share similar immune activation profiles (Geluk et al. 1993). In this context, it is evident that rhesus macaques represent excellent models to study various aspects of adaptive immunology in relation to human biology and disease. The diversity at the *MHC* class II region carries potential implications for experimental outcomes in biomedical research. For example, homozygosity of a specific *Mamu-DQ-DRB* haplotype (*DQB1*06:01* – *DRB1*03:09* – *DRB*W002:01*; #4 in figure 5) has been associated with an accelerated disease progression in SIV-infected macaques in comparison to animals without this allele combination (Sauermann et al. 2000). This observation aligns with an advantage of *MHC* heterozygosity in SIV-infected Mauritian cynomolgus macaques (O’Connor et al. 2010). Another study demonstrated reduced CD4^+^ T cell counts in SHIV-infected cynomolgus macaques that exhibit a particular *MHC* class II haplotype (M2), containing two functional *DRB* genes (Borsetti et al. 2012; Wiseman et al. 2013). Similarly, correlations of *HLA*-*DRB* alleles and haplotypes with resistance or susceptibility to HIV-1 infection have been documented as well (Malhotra et al. 2001; Lacap et al. 2008). In addition, associations with *HLA* class II haplotypes have also been reported for other infectious diseases and autoimmune disorders, such as tuberculosis and rheumatoid arthritis (Singh et al. 1983; Mangalam et al. 2013; Houtman et al. 2021; Martin et al. 2021). Several of these diseases are modelled in macaque species (Dijkman et al. 2019; Na et al. 2020). Collectively, association studies seem to underscore the direct impact of the *MHC* class II diversity on immune responses in the context of various diseases. Our haplotype characterization strategy offers an improved methodology to examine the cooperativity between *MHC* class II alleles, and in particular the number, combination and type of *DRB* copies, that coexist on the same haplotype. This additional information is key to better interpretate experimental outcomes in animal models, but also in human disease association studies.

Our comparative study demonstrates not only diversity in the *DR* region, but also reveals allelic polymorphism in the genes encoding the molecules that are involved in peptide loading. Two important members of this loading pathway are encoded by the *TAP1* and *TAP2* genes, the products of which load peptides onto MHC class I molecules. The influence of *TAP* variation on peptide loading remains ambiguous, although associations have linked polymorphisms to the susceptibility to several diseases (Ozbas-Gerceker et al. 2013; Meng et al. 2018; Praest et al. 2018). Our findings may help to further examine these disease associations, by exploring a potential long-range linkage disequilibrium in which *TAP* variants may impact the expression of the highly diverse *MHC* class I genes in macaques. Such a phenomenon has been encountered in chickens (Walker et al. 2011). In a similar way, the variation demonstrated for the genes encoding the DM and DO dimers might directly affect the loading of MHC class II molecules. The DM-dimer is able to transiently bind to the classical DR heterodimer, thereby facilitating peptide release and mediating the loading of high-affinity peptides (Anders et al. 2011). The HLA-DO dimer binds DM molecules at high affinity, which may modulate the DM-catalysed exchange of peptides (Guce et al. 2013). A limited variation has been documented for the *HLA-DM* and -*DO* genes, which potentially impacts antigen presentation and T cell selection (Álvaro-Benito et al. 2015; Alvaro-Benito et al. 2016). The impact of DR allotypes on different diseases is evident, but the putative contribution of *DM* and *DO* polymorphisms has not been studied conclusively. With our methodology, the combinatorial effect of all *MHC* class II loci, including *DM* and *DO*, might be better assessed in health and disease.

In conclusion, we demonstrate a long-read ONT sequencing approach utilizing adaptive sampling to resolve a complex immune region. This cost-and time-efficient strategy also allow to reassess the annotation and function of pseudogenes, which are often neglected in current genome studies. Here, we hypothesize that a pseudogene, *DRB9*, may contribute to the stabilization of an important genetic region that encodes the DRA molecule. On the contrary, the diversification of the macaque *DRB* region appears to be propelled by the inactive *DRB6* gene, which harbours two strong LTRs. This pseudogene potentially serves as a hotspot for homologous chromosomal recombination events, which reshuffles cassettes of paralogous genes, generating a wide spectrum of region configurations. These fundamental insights advance the refinement of rhesus macaque models that are eminent to study health and disease.

## Material and Methods

### Cells and genomic DNA extraction

Rhesus macaques of mostly Indian origin are housed at the Biomedical Primate Research Centre (BPRC) in an outbred breeding colony. Each animal has previously been characterized for its *MHC* class I and class II gene content by typing microsatellite markers. For this genomic characterization project, we selected 16 rhesus macaques, which represents the 21 most common *MHC* class II haplotypes in our colony. From these animals PBMCs were isolated from heparin whole blood samples that were obtained during annual health checks. Ultra-high molecular weight (UHMW) DNA was isolated from PBMC samples (± 6 x 10^6^ cells) of 12 animals using the Circulomics Nanobind CBB Big DNA Kit (Circulomics, NB-900-001-01) and following the manufacturer’s instructions. For comparison of DNA isolation techniques, HMW DNA was also isolated from six PBMC samples (± 1 x 10^6^ cells) using the Monarch® HMW DNA Extraction Kit for Cells & Blood (NEB, #T3050L) following the manufacturer’s instructions. Fragments smaller than 40 kb were depleted from these samples using the SRE XL kit (PacBio, SKU102-208-400) according to the manufacturer’s instructions. No differences in fragment length or yield were noted for the different DNA isolation kits. To dissolve the viscous (U)HMW DNA, the samples were heated to 70 °C for 5 minutes, followed by overnight incubation at room temperature. If needed, UHMW DNA samples were left to rest at room temperature for up to 72 hours or needle shearing (26G) was performed till the sample was homogenous. The concentration and purity of the gDNA isolates were determined using a Nanodrop and a Qubit platform. The latter method was performed in triple to ensure an accurate concentration measurement of the highly viscous samples. The fragment length was assessed for the first five isolated UHMW DNA samples by utilizing pulsed-field gel electrophoresis (PFGE) using a CHEF Mapper XA system (Bio-Rad) and the following settings: 0.5X TBE buffer cooled to 14 °C, initial switch time of 45s with a linear ramp to 145s, pulse angle of 120°, voltage gradient of 6 V/cm, and a total run time of 20 hours.

### Library preparation for long-read sequencing on an ONT GridION device

Sequencing libraries were prepared using two versions of the ONT Ultra-long DNA sequencing kit (SQK-ULK001 and SQK-ULK114), as the new V14 chemistry was released during the time of our experiments. In brief, UHMW DNA samples were dissolved in 750 ul elution buffer that was provided in the DNA extraction kits. Subsequently, the DNA fragments were labeled with sequencing adaptors in a tagmentation reaction, followed by a precipitation clean-up that involved an overnight incubation to elute DNA. Wide-bore pipet tips were used during library preparation to avoid DNA shearing. The final DNA library was loaded on R9.4.1 or R10.4.1 flow cells, depending on the used library preparation kit, SQK-ULK001 and SQK-ULK114, respectively. Remaining volumes of DNA library were stored at 4°C. After 4-8 hours, the flow cells were washed with the Flow cell Wash kit (EXP-WSH004, ONT) according to the manufacturer’s instructions. Washed flow cells were again primed and subsequently reloaded with remaining DNA library. Each flow cell had a sequencing time of approximately 48 hours with a MUX scan every 4-6 hours. In total, 2 to 4 flow cells were used per animal.

### Adaptive sampling and sequencing data

No target enrichment strategies, such as hybridization probes or guided Cas9 nuclease complexes, were applied during library preparation. Instead, we used adaptive sampling, a computational tool to enrich for sequences of interest. The real-time basecalling of DNA fragments that are pulled through the pores allows direct alignment of sequences to a reference library. Sequencing is only continued for the DNA fragments that show similarity to the library and thereby the region of interest is enriched in the sequencing output.

Our library contained chromosome 4 from the rhesus macaque reference genome (Mmul_10), which harbors the complete *MHC* cluster. In addition, we also enriched for the *KIR* gene region by adding chromosome 19 to the library. For some sequencing runs, the library was extended with chromosomes 7 and 13, targeting the complex regions encoding the B cell receptor. In this paper, we will only discuss the results that were generated for the *MHC* class II region. The generated reads were filtered for quality (q-score > 7) and minimum length (> 3 kb). The average amount of on-target data that was generated per flow cell was 3,1 Gb and the reads displayed an overall q-score of above 10 and a N50 length of 67,2 kb.

### Assembly of MHC class II clusters

The reads targeting the *MHC* class II cluster were filtered out from the total set of data by mapping all ONT reads to a reference exon library containing sequences from the IPD-MHC database and our previous transcriptome studies using Minimap2. On average, 140 reads that were at least 10 kB in length mapped to the *MHC* class II region. Overlaps of the reads were determined by sequence alignments, and subsequently a framework of the entire *MHC class II* region was manually constructed based on sequence similarities and SNPs. This framework represents a manual assembly of a haplotype, using large overlapping reads. All on-target reads were then mapped to the framework sequence using Minimap2 (version 2.26) to confirm the region assembly and to generate a consensus sequence (Li 2018). For four haplotypes the central *MHC class II* region (*Mamu-DOB* to -*DOA*) could not be completely determined due to low coverage (< 3X) (Suppl. Table S1). The assembled regions were validated by cross-referencing with haplotype information on the classical *MHC class II* genes, obtained from STR typing and segregation data, which is available for all rhesus macaques housed at the BPRC.

### High coverage sequencing

Phased *MHC* class II regions of one heterozygous rhesus macaque were characterized at high accuracy to determine gene sequence similarities compared the human reference genome. For this animal, HMW genomic DNA was extracted as described above. Subsequently, an ONT and PacBio hybrid sequencing approach was performed. This involved sequencing on a R10.4 Promethion flow cell using the Ligation Sequencing Kit V14 (SQK-LSK114) and an ONT P2Solo device in combination with HiFi sequencing on a SMRT cell using the SMRTbell® prep kit 3.0 and a PacBio Revio device. In addition, two other samples (R02034, R04022) were sequenced using a Promethion flow cell and the ONT P2Solo device as described above, without complementation of PacBio sequencing or utilization of adaptive sampling.

The PacBio and/or ONT reads obtained for these three samples were filtered on length (> 5kb) and quality (q-score > 10) before they were mapped to the reference sequence (Mmul_10) using Minimap2. All reads that mapped to the *MHC class II* region were extracted. The combination of PacBio and ONT reads was assembled using Hifiasm (version 0.19.8) with default settings and ultra-long ONT integration, which produced contiguous phased regions with over 30X coverage (Cheng et al. 2021). The phasing of the assemblies was validated using STR typing and segregation information of the classical *MHC class II* genes that was available for this animal. The ONT reads from the other two samples were assembled using a similar method as described above for the lower coverage haplotypes.

### Annotation, similarity and pylogenetic analysis, and visualization

The assembled *MHC class II* regions were annotated based on sequence similarity to the reference sequences derived from the IPD-MHC (Release 3.11.0.0) or reference genome (Mmul_10). Similarity was determined by alignment of the exon reference sequences to the assembled consensuses using Minimap2. Genes displaying variation in their exons, as compared to the reference sequences, were designated based on the nearest reference match, augmented by the suffix “new” (e.g., *Mamu-DMB*03:02new*). When multiple novel sequences shared the closest reference match, the suffix was accompanied by sequential numbering (e.g., “new2”). Intronic variations identified for gene copies present on the different assembled *MHC* class II regions were assigned temporary names in accordance to the standardized nomenclature system for non-human primates (de Groot et al. 2012).

The annotated macaque genes were extracted from the assemblies as BED files, which were then converted to FASTA format using bedtools (v2.26.0). These FASTA files containing the macaque sequences were aligned against the human reference genes using blastn (v2.6.0+) to determine sequence similarity. Genes were considered homologs when they shared at least 85% similarity. Only the blast hits with the highest scores and with an alignment length of over 1000 bp were retained. Package gggenomes was used to generate a visualization of the macaque *MHC* class II assemblies in alignment to human reference sequences, thereby indicating the homology of the annotated genes.

Neighbour-joining trees were generated for exon 2 sequences of the *DRB* genes using Geneious prime software applying the Jukes-Cantor and the Kimura substitution models. Both approaches demonstrated separate clusters for *DRB6* and *DRB9* sequences, substantiated by high bootstrap values.

## Data availability

All raw and processed sequencing data generated in this study has been submitted to the European Nucleotide Archive (ENA, https://www.ebi.ac.uk/ena/browser/home) under project number PRJEB70961. The annotated consensus sequences of the *MHC* class II haplotypes have been submitted under accession numbers ERZ23795125 to ERZ23795148.

## Acknowledgements

We thank Francisca van Hassel for design of figures and artwork.

## Author information

These authors share last authorship: Jesse Bruijnesteijn, Ronald E. Bontrop

## Contributions

J.B., N.G.G., R.E.B. conceived and designed the methodology. N.G., M.W., N.G.L. and J.B. performed the experiments and N.G., J.B., N.G.L., N.G.G., R.E.B. analysed the results. All authors discussed results and interpreted data. J.B. and R.E.B. co-wrote the paper. All authors approved the submitted version of the manuscript.

